# Immunological Patterns from Four Melioidosis Cases: Constant and Variable Protein Antigens

**DOI:** 10.1101/082057

**Authors:** Jinhee Yi, Kelsey Herring, Timothy C. Sanchez, Srinivas Iyer, Joshua K. Stone, Judy Lee, Mark Mayo, Bart J. Currie, Apichai Tuanyok, Paul Keim

## Abstract

*Burkholderia pseudomallei* is the causative agent of the melioidosis and is endemic to Southeast Asia and northern Australia. There is no available vaccine and accurate diagnosis is difficult, time-consuming and labor intensive. Early diagnosis is an important part of successful treatment and current serological tests are inadequate and based upon multiple antigens. Identifying specific immunogenic proteins which are highly seroreactive may yield potential diagnostic targets for detecting antibodies and antigens specific to melioidosis. We have used 2D gel electrophoresis and Western blotting analysis to analyze protein antigenicity of whole cell lysates extracted from four *B. pseudomallei* strains and the sera from the specific infected humans. We found a total of 135 immunogenic proteins, 62 of which we were able to identify to a specific gene by mass-spectrometry. Results from the Western blotting of each strain’s proteins and the corresponding patient serum reveal between 30 – 40% serum x strain specific immunogenic proteins. In most cases, these differences exist despite the fact that the genes encoding these proteins were present among all four *B. pseudomallei* strains. Eight particular proteins were immunogenic in all four strain x serum combinations and could represent novel diagnostic and vaccine subunit targets.

## Introduction

*Burkholderia pseudomallei* is a Gram-negative environmental saprophyte and is also the causative agent of melioidosis, a potentially deadly disease endemic to Southeast Asia and northern Australia [1]. It is now being recognized in more diverse global regions such as South America, the Caribbean and Africa [1–3]. Melioidosis can manifest in numerous clinical forms, most commonly as pneumonia, but also a spectrum from localized cutaneous disease without sepsis to rapidly progressive fatal septicemia [1, 4]. Risk factors have been identified which contribute to the likelihood of developing melioidosis. Diabetes mellitus, hazardous alcohol use, renal disease, chronic pulmonary disease, immunosuppressive therapy and thalassemia are all associated with an increased risk for melioidosis. The mechanism by which these risk factors impact the disease is not entirely clear, but having any of them can result in more severe disease with higher mortality [1, 5].

First-line treatment for melioidosis utilizes β-lactam antibiotics such as ceftazidime, but *B. pseudomallei* can develop resistance to these antibiotics during the course of acute infection and eradication phases [6, 7]. This acquired resistance may be linked to efflux pumps, enzymatic inactivation, and alteration of drug targets and decreased permeability [8].

*B. pseudomallei* is a resilient bacterium that can tolerate hostile conditions by producing and secreting proteases, lipases, catalases, peroxidase and siderophores. In addition, it can evade host immune responses and is able to survive within phagocytic cells [9]. *B. pseudomallei* produces virulence effectors that can be transferred into host cells through different machineries. For instances, the type III and VI secretion systems have been identified as allowing the bacteria to survive intracellularly, escape autophagy and spread within the host [10–12].

Surface proteins of this organism have received attention as the majority of protein antigens that are recognized by host response antibodies [8]. Such surface antigens are critical targets of humoral immune response and have the potential to be developed for immunological diagnosis and therapies [9]. However, to facilitate the development of better diagnostics, comprehensive information of these antigens is needed. Identifying such proteins in *B. pseudomallei* can form the basis for all these purposes. In this study we determined the immunogenic protein profiles in four *B. pseudomallei* strains which were isolated from four different human melioidosis cases using 2D gel electrophoresis, Western blot hybridization, and mass-spectrometry. The immunogenic protein profiles for all four strains were determined using matched sera from the infected patients.

## Materials and Methods

### Ethics Statement

Our study examined two primary B. pseudomallei isolates (MSHR) obtained from 556 melioidosis patient group from the Darwin Prospective Melioidosis Study, which commenced at Royal Darwin Hospital, Northern Territory, Australia in October 1989. Ethics approval for this study was obtained through the Human Research Ethics Committee of the Northern Territory Department of Health and Menzies School of Health Research, approval number HREC 02/38 (Clinical and Epidemiological Features of Melioidosis) with written informed consent obtained from patients. All patient data were de-identified prior to analysis. The two Arizona USA patients’ specimens (PB) were collected for clinical diagnostic purposes and deidentified prior to transfer to NAU. As such, they are exempt from USA IRB regulations.

### Bacterial Strains and Growth Conditions

Four human clinical *B. pseudomallei* strains MSHR1079, MSHR1328, PB08298010, and PB1007001 were used in this study. Three strains were isolated from blood culture - negative localized infection (e.g., skin, lungs), while the other one was from a blood culture positive case. Details of these strains, sera and clinical information are shown in Table 1. We used Luria-4 Bertani (LB) agar to grow the bacterial strains at 37°C for 48 hours subjected to protein extraction.

**Table 1.** Summary of *Burkholderia pseudomallei* strains and human patient serum used in this study.

| Strain/ LPS genotype | Country of origin | Sera and Clinical information |
| --- | --- | --- |
| MSHR1079/ serotype B | Australia | This was a 50 year old patient who was sick for 6 weeks before the diagnosis of blood culture-negative skin infection by culture of *B. pseudomallei* from the regional lymph node. Clinical manifestation was an arm abscess with lymph node involvement. IHA titer was very high on admission (1:5120). The serum was collected from when the treatment was started, i.e., at the time of initial diagnosis. The patient was treated initially with intravenous ceftazidime plus oral trimethoprim/sulfamethoxazole and made a full recovery (Tuanyok et al.[15]). |
| MSHR1328/ serotype A | Australia | This was a 29 year old diabetic who was sick for 2 weeks with mild symptoms before the diagnosis of blood culture-negative pneumonia. Only sputum was culture positive. IHA titre was very high on admission (>1:5120). The serum was collected from when treatment was started, i.e., at the time of initial diagnosis. The patient was treated initially with ceftazidime and made a full recovery (Tuanyok et al. [15]). |
| PB08298010/ serotype A | Arizona, USA | This was a 32 year old man who was sick for 6 weeks with a history of type II diabetes, hypertension, and obesity. Synovial fluid, blood and sputum were culture positive. Clinical manifestations were severe right knee pain and febrile upon hospital admission but the patient's hospital course was complicated by respiratory failure that required intubation and ventilation, acute renal failure, pneumothorax and pneumoperitoneum, anemia and hypotension. Antibiotics treatment included with meropenem, moxifloxacin, vancomycin, ceftazidime, gentamicin, and trimethoprim/sulfamethoxazole. All cultures were negative 2 weeks before discharged from hospital. The serum was collected 7 weeks after discharged from hospital. Source of exposure is unknown (Stewart et al. [16]). |
| PB1007001/ serotype A | Arizona, USA | This was a 49 year old woman who was sick for 8 weeks with a leg skin abscess and a swollen regional lymph node. Skin abscess fluid was culture positive for *B. pseudomallei*. Before diagnosis treatment included clindamycin, imipenem, vancomycin and metronidazole. The patient was treated with intravenous ceftazidime and oral trimethoprim/sulfamethoxazole. The time of serum collection is unknown. The epidemiologic investigation revealed that the patient had traveled to Costa Rica three months before the patient developed symptoms, suggesting infection acquired overseas. |

### Protein Extraction

For each protein preparation, bacterial colonies were suspended in sterile 1X PBS, pH 7.3, until a turbidity of 1.5 at OD_600_ was obtained. The bacterial cells were pelleted by centrifugation at 16,000 × *g* for 3 minutes. The supernatant was removed and the cell pellet was resuspended in 500 µL of lysis buffer (0.5% Triton X-100; 50 mM potassium phosphate, pH7.8; 400 mM NaCl; 100 mM KCl; and 10 mM imidazole). The solution was subjected to 3 freeze & thaw cycles alternately in liquid nitrogen and a 42°C dry bath to lyse the cells. The sample was then centrifuged at 18,000 × *g* for 15 minutes at 4°C to separate the soluble (liquid) from the insoluble (pellet) proteins

The quantity of each protein sample was determined using the Bradford assay with slight modifications [17]. Bovine serum albumin (Bio-Rad, Hercules, CA, USA) was used as a standard protein. Prior to protein quantification, the sample was rinsed and equilibrated in Tris-HCl buffer.

### Two-dimensional electrophoresis (2DE)

First dimensional isoelectric focusing (IEF) and second dimensional SDS-PAGE were performed as previously described [18–20]. The IEF was conducted in the pH 4-7 range after exploring a wider range (pH 3-10) that provided little additional resolution. Most proteins were found to have a pI value between 4.5 and 6.5 and a molecular weight of 20 to 100 kDa. Each protein sample was treated with trichloroacetic acid (TCA) and acetone for purification. A total of 100 µg of each protein sample was tested on each 2DE. The pellet was resuspended in 160 µL of a resuspension buffer.

The IEF was performed using 2D Electrophoresis ZOOM® IPGRunner System (LifeTech, Carlsbad, CA, USA). Briefly, a 7cm IPG strip, pH 4–7, was loaded with 160 µL of the protein sample, and then operated at 6,000 VHr. The IPG strip was treated with the reduction buffer (130 mM DTT, 5 M urea, and 0.8 M thiourea).for 20 minutes and with an alkylation buffer (130 mM iodoacetamide, 0.002% bromophenol blue, 5 M urea, and 0.8 M thiourea). Proteins immobilized on the IPG strips were further separated using SDS-PAGE with a 4-20% Tris-glycine gradient gel (ZOOM® Novex gel, LifeTech). Electrophoresis was performed at 110 V for 90 min, and the gel was then visualized by silver staining according to Shevchenko’s method [21]. Gel image was digitalized by UVP gel documentation system and analyzed using the VisionWork LS (UVP, Upland, CA, USA).

### Western blotting analysis

A duplicated non-silver stained 2D gel was blotted transferred to a nitrocellulose membrane in a dry transfer system (iBlot® Dry Blotting system, LifeTech) for 10 minutes at 25mA. The blotted membrane was blocked with 2.5% skim milk in 1X PBS for 1 hour. The membrane was incubated with the patient serum solution (1:1,000 v/v in 2.5% skim milk) for 1 hour and then washed in 1X PBS three times. The membrane was incubated with a 1:1,000 dilution of HRP-conjugated goat anti-human IgG secondary antibody (Promega, Madison, WI, USA) in 2.5% skim milk followed by an additional three washes in PBS. Then, the immunogenic protein spots were developed colorimetrically using 3,3'-Diaminobenzidine solution.

### In-Gel Trypsin Digest

The immunogenic protein spots on the Western blot membrane were matched with 2D silver stained gel image using 2D analysis software (Melanie v. 7.0.6, Genebio, Geneva, CH). The matched immunogenic spots were excised from the gels and destained with 0.02% sodium thiosulphate and 0.5% potassium ferricyanide based on the method of Shevchenko *et al.,* (1996). Excised gel pieces were washed and dried with 50% acetonitrile and then reduced and alkylated in 10 mM DTT and 100 mM iodoacetamide. Proteins were digested in-gel overnight with 12.5 ng/mL trypsin (Promega Inc., USA) made up in digestion buffer (50 mM ammonium bicarbonate; 5 mM calcium chloride) at 37°C. The cleaved peptides were extracted using extraction buffer made up of 5% formic acid and 50% acetonitrile. The extraction of digested peptide was facilitated by vortexing for 30 minutes and sonication for 20 minutes. The extracts were placed in new tubes and dried completely [21, 22].

### Matrix Assisted Laser Desorption/Ionization Time-of-Flight Mass Spectrometry (MALDI-TOF MS)

The dried peptide samples were reconstituted in 4µL of 0.1% Trifluoroacetic acid (TFA) in water. Samples were mixed 3:1 ratio (matrix:sample) with 5 mg/mL of Ultrapure α-cyano-4-hydroxycinnamic acid (CHCA) matrix (Protea, Morgantown, West Virginia). Prior to sample analysis, the mass spectrometer was externally calibrated with a TOF/TOF Calibration peptide mixture of des-Arg-Bradykinin (1.0 pmol/µL), Angiotensin I (2.0 pmol/ul, Glu-Fibrinopeptide B (1.3 pmol/µL), and adrenocorticotropic hormone (ACTH), (1-17 clip-2.0 pmol/ µL), (18-39 clip-1.5pmol/ µL), (7-38 clip-3.0 pmol/ µL). The 4800 Plus MALDI TOF/TOF Analyzer (ABSciex, Foster City, CA, USA) was used in positive reflector mode. Data were collected in an automated Batch mode utilizing random sampling over the entire sample spot. The mass spectrometer is equipped with a 200-Hz frequency Nd:YAG laser, operating at a wavelength of 355 nm. Twenty-five sub-spectra for each of 25 randomized positions within the spot (625 spectra/spot) were collected in MS-TOF mode and presented as one main spectrum. MS/MS, fragmentation mode, 50 sub-spectra for each of 13 randomized positions within the spot (650 spectra/spot). Fixed laser intensity in MS-TOF reflector positive mode, 3600 (arbitrary units), final detector voltage set at 1.883 KV. MS/MS-TOF/TOF, Fragmentation mode, fixed laser intensity set at 4500 (Arbitrary units), final detector voltage set at 2.080 KV. Spectral Mass range 850-4000 m/z, focus mass 2250 m/z. The MS/MS data were analyzed using the Paragon Algorithm [23] of ProteinPilot Software version 4.0 with settings: *Sample type*: Identification; *Cys Alkylation*: Iodoacetamide; *Digestion*: Trypsin; *Instrument*: 4800; *Species*: (no filter applied); *Special factors:* Gel-based ID; *Search Effort*: Thorough; *ID focus*: none applied; *Database*: Specific strain-translated genome database; and the *Detected Protein Threshold*: (10% confidence). Peptide mass and fragmentation spectra were searched against a protein database which was generated from each strain by whole genome sequencing and RAST annotation [24].

### Bioinformatic analysis of immunogenic proteins

Identification of immunogenic proteins was accomplished by determining peptide sequences with MS/MS and mapping them to known sequences generated by RAST and NCBI’s databases. After identifying antigenic proteins, the observed pI and molecular weight values were compared to the predicted values of each to assess the proper identification of proteins. Predictions of cellular location and function (including signal peptides) were utilized to determine if any identified outer membrane or surface proteins were putative virulence factors. FIGfam classification was used to categorize and determine the functional group of each immunogenic protein [25]. The majority of immunogenic proteins were proteins known or predicted to be involved in various metabolisms (see Table 2).

### Protein Identification

For proteomic information on identified immunogenic proteins, several web-based tools were used. Cellular location of identified proteins was predicted using the program “PSORTdb” version 2.0 [26]. The presence of signal peptides was inferred using “SignalP” version 4.0 [27]. This program was used to hypothesize N-terminal secretory signal peptides of the identified immunogenic proteins. Proteins with a SignalP D-score > 0.57 were considered to be potentially translocated. Protein identity was analyzed using BLASTP against our RAST-generated database. Theoretical molecular weight and pI values were calculated using Compute I/Mw tool on ExPASy website (http://web.expasy.org/compute_pi/).

## Results

### Experimental design and serum selection

Because of the great genomic diversity among *B. pseudomallei* strains [28], we used matched sets of human sera and protein extracts of isolates from these same melioidosis patient (Table 1). This was done to insure that at least the genes for antigenic proteins are present with the potential to express and to be detected by the host. In one preliminary experiment, we did examine the antigenicity of proteins expressed *in vitro* from a single strain (PB08298010) against sera from its source patient but also two others (PB08298010 and MSHR1328). We found that the two heterologous cross-reaction Western blots detected fewer immunogenic proteins than its source patient’s serum (data not presented). This is consistent with differential strain gene 11 contents, but it should also be noted that each melioidosis case is unique with a different host, infection time sequences and complex pathologies that could alter antigen-humoral responses.

## Immunogenic protein profiles

A total of 135 protein spots were confirmed to be immunogenic across all four melioidosis cases (Fig 1). The cross reactivity of these 135 proteins was distributed in a complex manner across the four infections (Fig 2). PB1007001 contained a greater proportion of strain X patient specific antigenic proteins (52%) than the other three strains (30 – 44%). The two Arizona cases had a higher number of total immunogenic proteins than the two Australian strains with the presence of seven common immunogenic proteins in all four infections.

Fifty unique proteins were identified using MS and ascribed to specific genes (Table 2). The 10 immunogenic proteins shared by all four melioidosis cases included the chaperone GroEL, elongation factor Tu (EF-Tu), enolase, alkyl hydroperoxide reductase protein C, NADH dehydrogenase, cell division protein FtsA,sigma-54 dependent DNA-binding response regulator and LysR-family transcriptional regulator (LTTR). GroEL and EF-Tu have each isoform commonly detected in all the four strains.

## Comparison of identified common immunogenic proteins

The eight common immunogenic proteins were BLASTed against the protein sequences of four human commensal bacteria and two genetically related strains of *Burkholderia mallei* and of *B. thailandensis* (Table 3). This sequence comparison was to evaluate the potential of the common immunogenic proteins to be used as a diagnostic biomarker or vaccine target candidate. For protein sequence BLAST, the sequence of *Staphylococcus aureus* NCTC8325, *Clostridium perfringens* F262, *Lactobacillus acidophilus* ATCC 4796, *Escherichia coli* MG1655, *Burkholderia mallei* ATCC 23344, and *B. thailandensis* E264 were used. The BLAST results showed there were low similarities with the four selected commensal bacteria. Most similarities were less than 50% and even the highest similarity of GroEL and EF-Tu showed only 76% and 80%, respectively, with the commensal bacteria. There are likely unique epitopes in the *B. pseudomallei* proteins that would not be confounded by infections by these or related bacteria. As would be expected, the sequence similarity of *B. mallei* and *B. thailandensis* was higher with an almost 100% similarity. However, Ndh and AtoC proteins of non-pathogenic *B. thailandensis* E264 showed a relatively lower similarity than *B. mallei* ATCC23344 as evidenced by 93% and 94% respectively. This illustrates the complications for diagnostics in distinguish infections by closely related bacteria.

**Figure 1:**
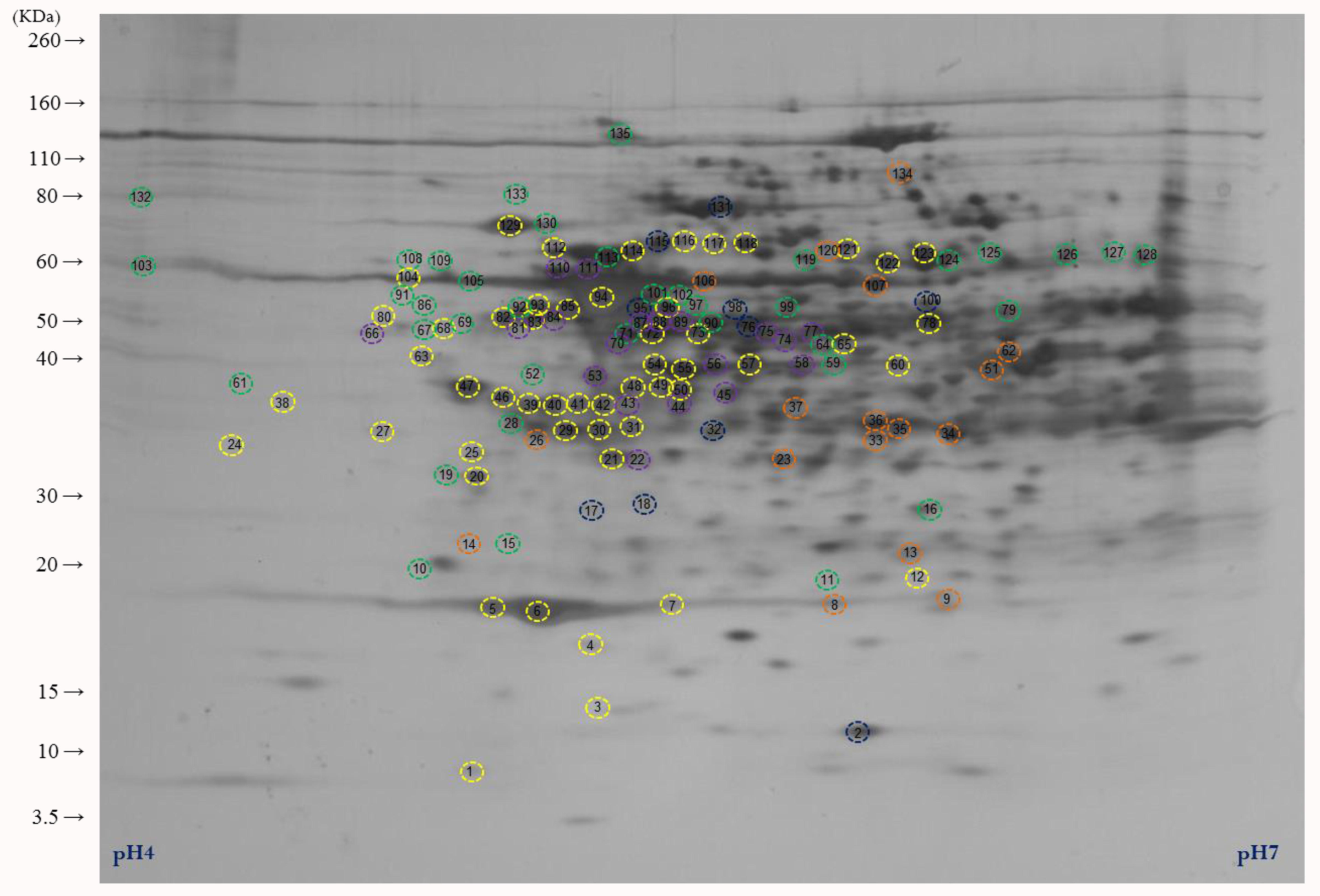
2D silver stained map of immunogenic proteins of four *Burkholderia pseudomallei*strains. Colored circles highlight all immunogenic proteins identified. Proteins identified in at least two strains as immunogenic are colored in yellow and in only one strain as follows: MSHR1079 (blue), MSHR1328 (orange), PB08298010 (purple), and PB1007001 (green).

**Figure 2:**
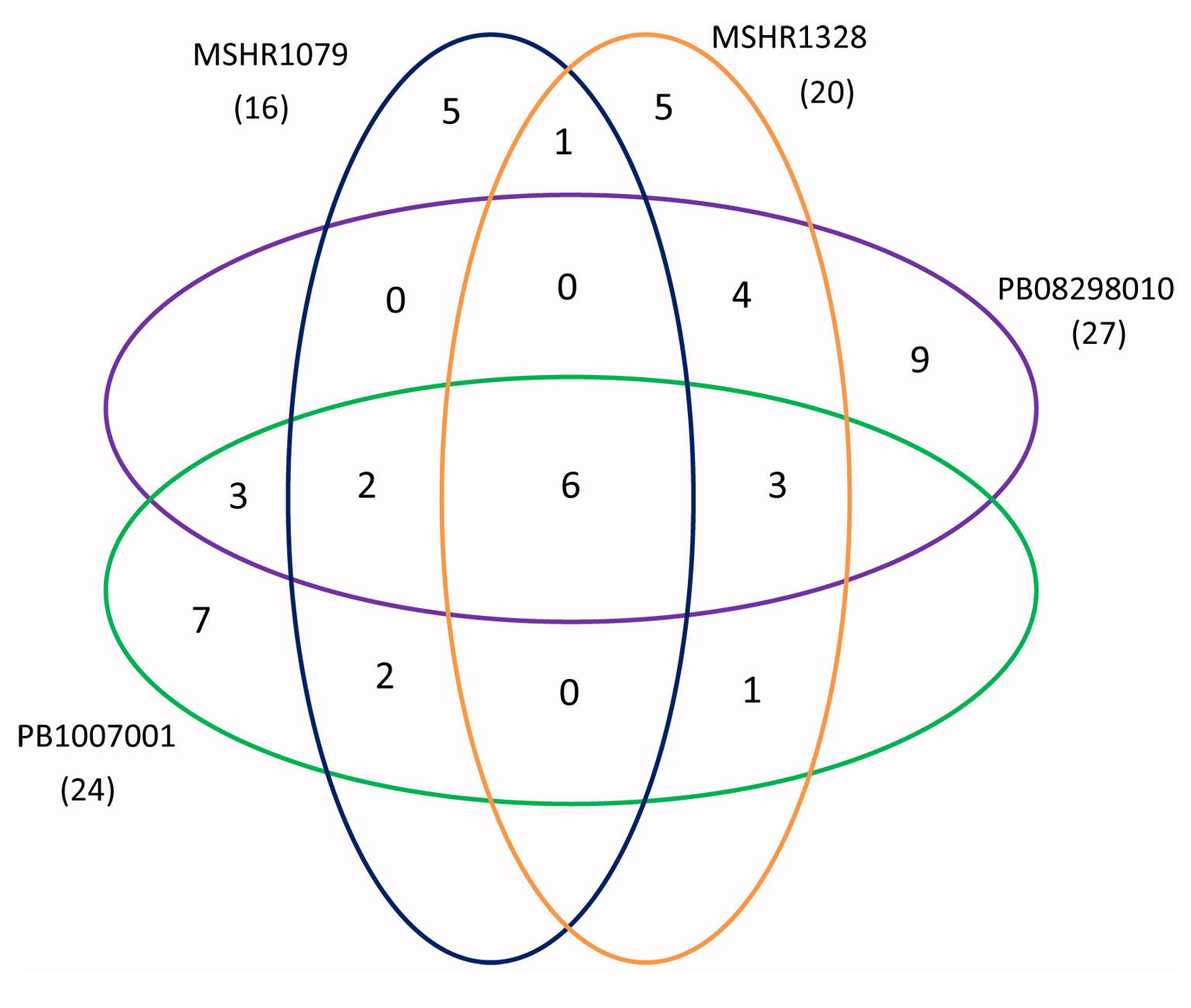
Comparison of immunogenic proteins identified from four *Burkholderia pseudomallei* strains. Number is the total detected immunogenic protein spots for each strain and numbers in parentheses are the number of identified immunogenic protein spots for each strain by mass spectrometry. Colored circles correspond to each strain as follows: MSHR1079 (blue), MSHR1328 (orange), PB08298010 (purple), and PB1007001 (green). Note: This 2DE image was generated from *B. pseudomallei* strain PB0829010.

**Table 2.**
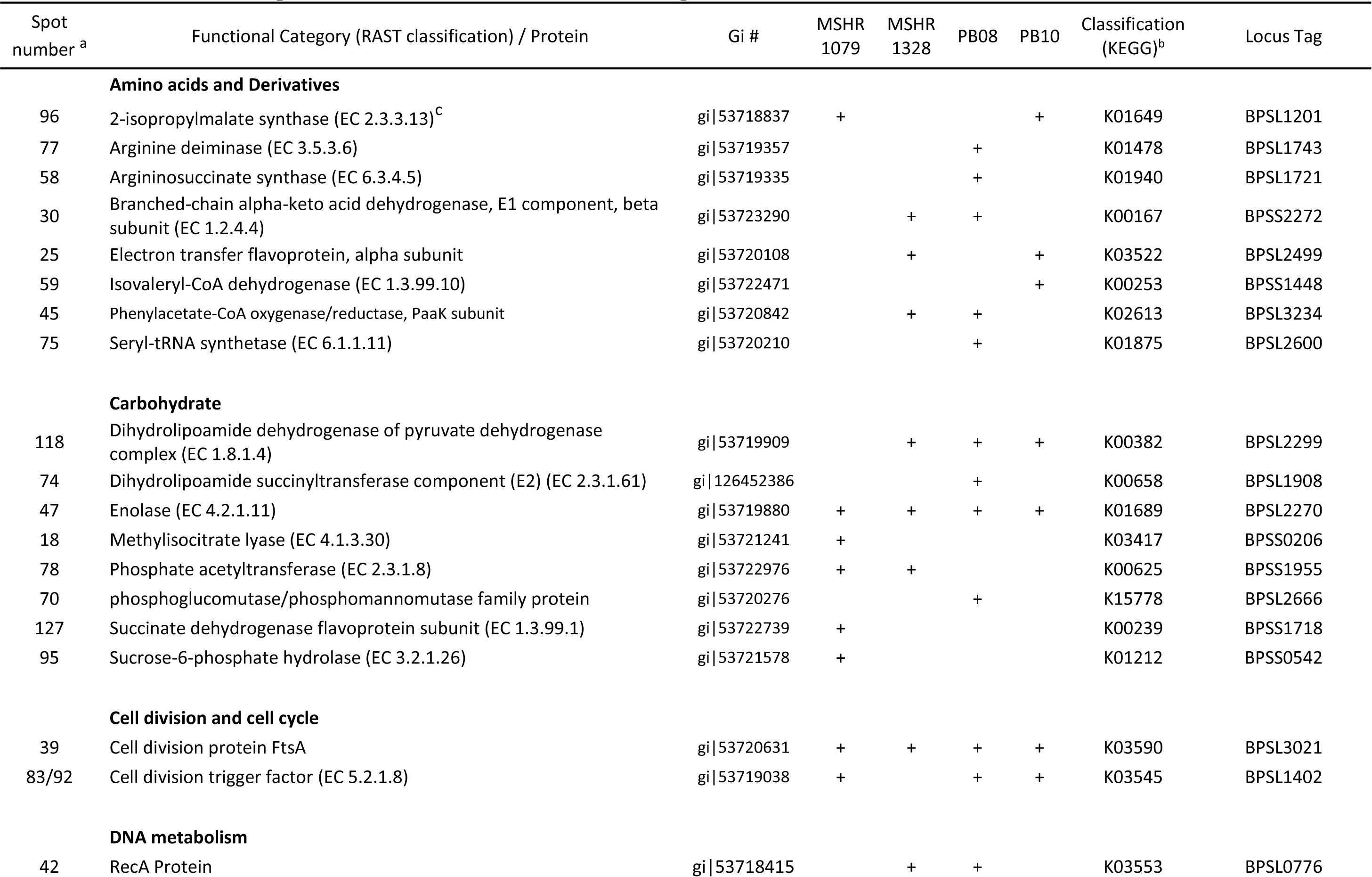
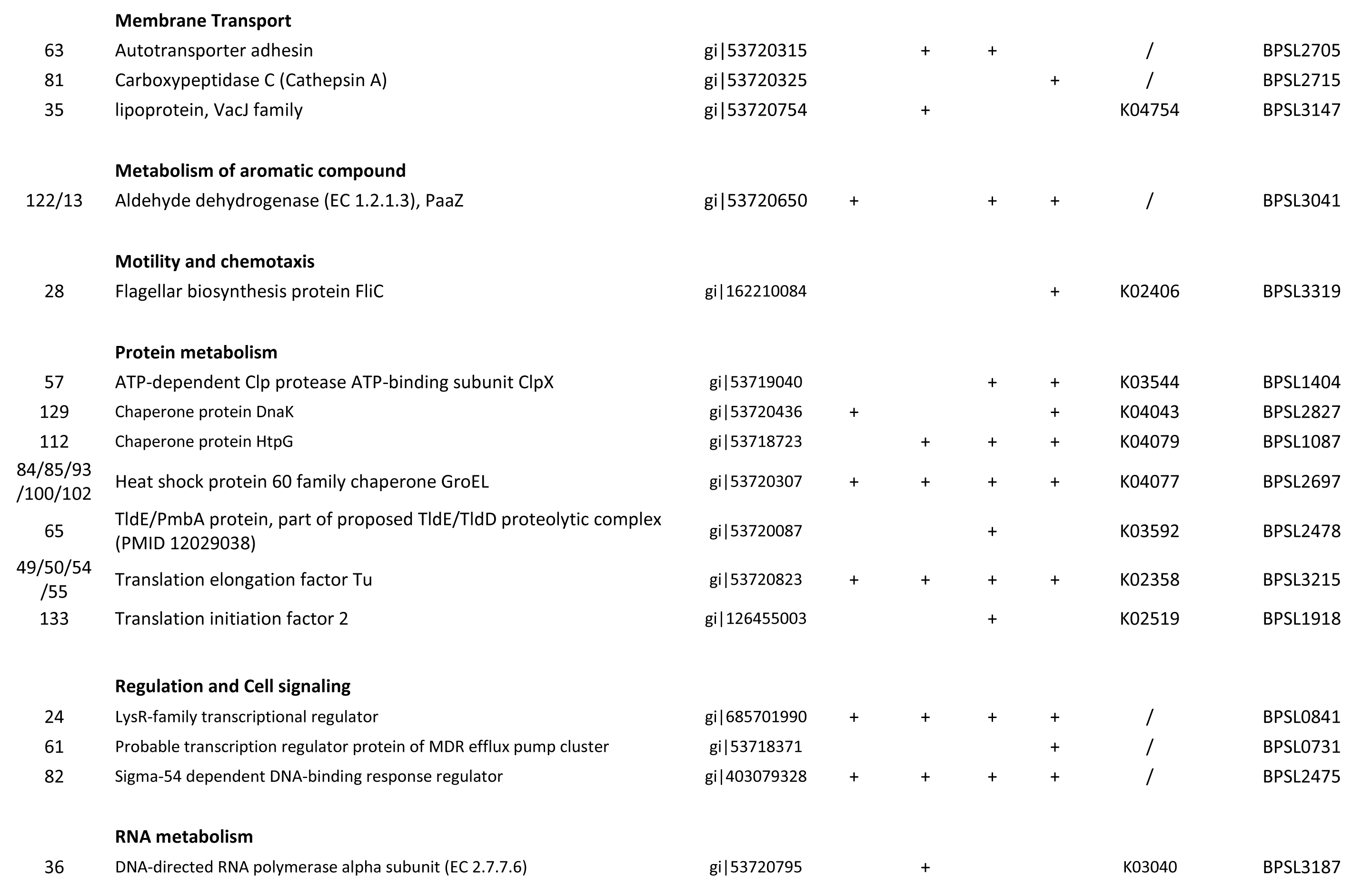
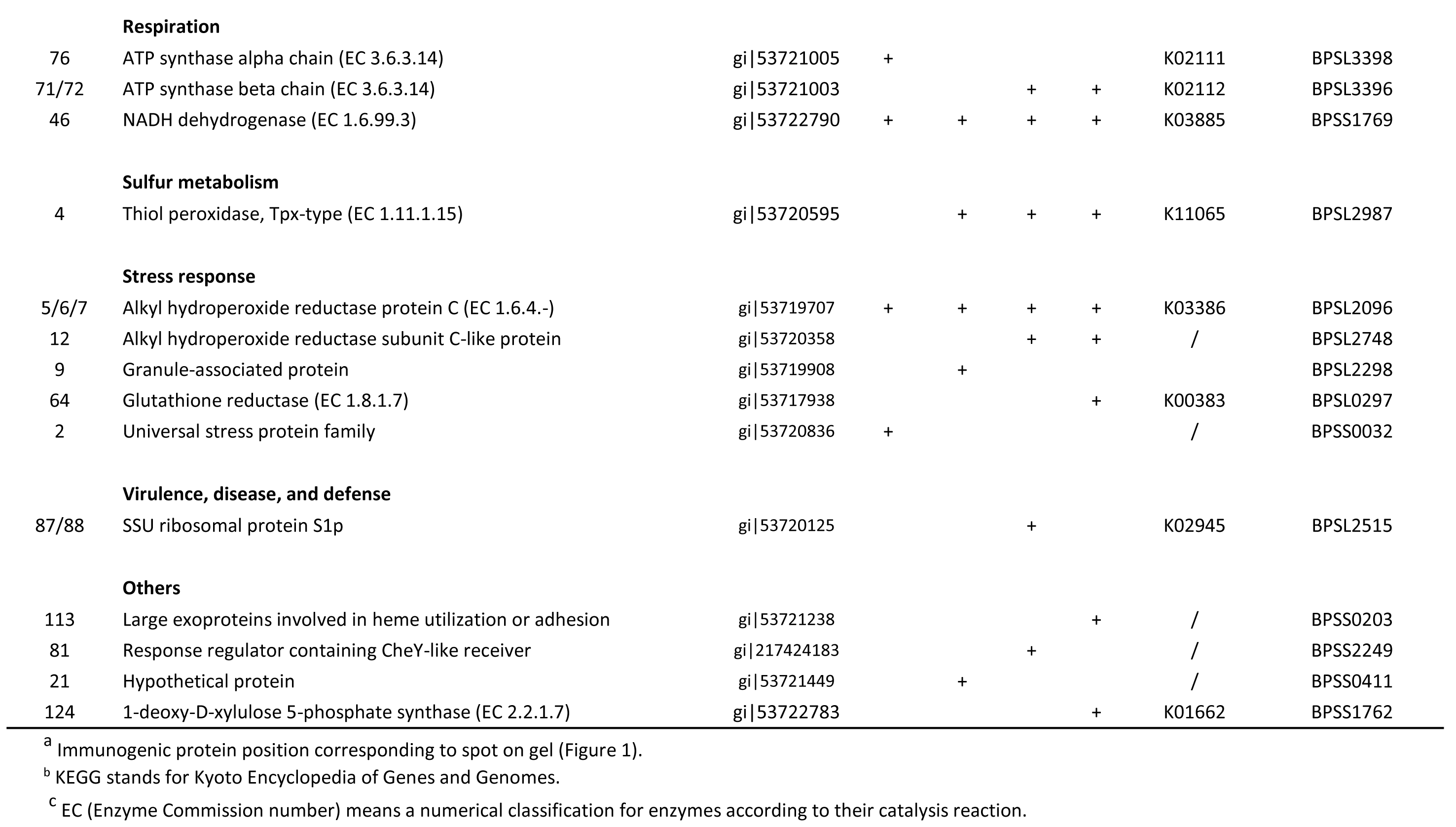
Immunoreactive proteins identified from four *Burkholderia pseudomallei* strains

**Table 3.** Percent identity of immunogenic proteins of *B. pseudomallei* compared to four commensal bacteria and *Burkholderia mallei* and *Burkholderia thailandensis* E264.

| Species | % Identity of *Burkholderia pseudomallei* antigenic proteins relative to other bacteria |  |  |  |  |  |  |  |
| --- | --- | --- | --- | --- | --- | --- | --- | --- |
|  | GroEL | EF-Tu | Eno | AhpC | Ndh | FtsA | AtoC | LTTR |
| *Staphylococcus aureus* NCTC8325 (taxid:93061) | 58% | 77% | 64% | 28% | 25% | NS | NS | 21% |
| *Clostridium perfringens* F262 (taxid:883064) | 60% | 75% | 65% | 32% | NS * | 27% | 41% | NS |
| *Lactobacillus acidophilus* ATCC 4796 (taxid:525306) | 58% | 26% | 51% | 33% | 24% | 26% | NS | NS |
| *Escherichia coli* MG1655 (taxid:511145) | 76% | 80% | 66% | 33% | 45% | 47% | 46% | 30% |
| *Burkholderia mallei* ATCC 23344 (taxid:243160) | 99% | 100% | 100% | 100% | 100% | 99% | 99% | 99% |
| *Burkholderia thailandensis* E264 (taxid:271848) | 99% | 100% | 99% | 100% | 93% | 99% | 94% | 99% |

\* NS: No Significant Similarity
(A) Heat shock protein 60 family chaperone, GroEL (B) Translation elongation factor Tu, EF-Tu (C) Enolase, Eno (D) Alkyl hydroperoxide reductase subunit C, AhpC (E) NADH dehydrogenase, Ndh (F) Cell division protein, FtsA (G) Sigma-54 dependent DNA binding response regulator, AtoC (H) LysR family transcriptional regulator, LTTR.

## Discussion

### Immunogenic protein profiles

Our findings demonstrate common and unique immunogenic proteins across the four *B. pseudomallei* strains from different melioidosis cases. Given the different clinical presentations and time courses of infection and disease for each serum collection, it is to be expected that the primary immunogenic proteins would vary across the matched four strain sera [29]. It has been postulated that variation in immunogenic proteins can be related to different stages of illness and the different clinical presentation’s related to particular virulence pathways.

We identified 50 immunogenic proteins from 135 total immunogenic protein spots by MS. Most proteins identified in this study had not been previously described as immunogenic in *B. pseudomallei*, but six had been identified as immunogenic outer membrane proteins by Harding *et al.* [30] and seven proteins detected in this study were found by Felgner *et al.* [14]. These proteins are GroEL, AtpD, AtpA, DnaK, EF-Tu, AphC, FliC, and sucrose-6-phosphate hydrolase. Most previous studies have used pooled sera and then investigated whole cell lysate samples, fractionated proteins, or preselected immunogenic recombinant protein candidates [14, 31–33]. We detected fewer immunogenic proteins when compared to other studies. Our use of whole cell lysates may be a limiting factor that prevents us from detecting the immunogenicity of low concentration proteins. However it also allows us to determine the relative protein abundance and identify highly immunogenic proteins. Analyzing each strain with only one serum time-point from each patient may restrict our ability to detect additional immunogenic proteins as the patterns may change as the disease progresses. There are eight common immunogenic proteins observed across all four strain x serum combinations.

### Chaperone GroEL

Of the identified immunogenic proteins, heat-shock proteins (hsps) were detected with great intensity and seroreactivity. Woo *et al.* [31] showed that sera from melioidosis patients showed a stronger antigen-antibody response to heat-shock protein 60 family chaperone GroEL than sera from patients infected with *Burkholderia cepacia* complex strains. Amemiya *et al.* [32] also found that GroEL was far more immunoreactive than DnaK. From this, it is not surprising that GroEL is a dominant immunogenic protein. Varga and colleagues [34] also found that the antibodies against GroEL increased almost 5 fold after infection [34]. *B. pseudomallei* has two copies of GroEL (BPSL2697 and BPSS0477) with 83% sequence identity. Only GroEL1 (BPSL2697) was detected in this study.

As a Hsp, GroEL is known to be involved in assisting protein folding *in vivo*. This folding process is subjected to interference from elevated temperatures, changing pH, and oxidative stress. Given the high conservation of hsps, particularly among related bacteria, it is possible that the host immune system is “primed” to target common epitopes of hsps, particularly GroEL, explaining its high seroreactivity with melioidosis. This idea is consistent with the work of Yamaguchi *et al.* [35], who found an epitope on *Helicobacter pylori* GroEL that was shared with numerous other species.

### Alkyl Hydroperoxide Reductase Protein C

Free radicals and oxidative stress play important roles in the innate immune response of the host. Phagocytic cells are able to kill invading bacteria through a pathway reliant on reactive oxygen and nitrogen intermediates [36]. Two of these roles are to inhibit proteases via S-nitrosylation of the cysteine residues in the activation site and nicking of DNA to induce apoptosis [36–39]. In this study, AhpC (Akyl hydroperoxide reductase protein C), a predicted oxidative stress response protein was detected in all four strains. As AhpC was highly expressed, it is possible that the expression of this protein is up-regulated during the early stages of infection and phagocytosis when oxidative stress response is at its greatest in the host [36, 40]. If this postulate is correct, then AhpC may represent an attractive target for early immunological diagnosis of melioidosis.

### Enolase

The main function of enolase is a part of the glycolysis pathway, but it has also been described as a “moonlighting” protein, detected on the cell surface and involved in plasminogen binding. This additional function allows the bacteria to penetrate the extracellular matrix and cell membrane, facilitating the spread of bacteria into tissue. Enolases do not have traditional sorting signals and it is unknown how the proteins are transferred to the cell surface [41–43].

The amino acid sequence of enolase in *B. pseudomallei* (BPSL2270) is rich in alanine, glycine, and leucine (11%, 10.8%, and 10.1%, respectively). Lysine is recognized as an important amino acid in the active site of the C-terminus and comprises 4.7% of the sequence in BPSL2270. A hydrophobic domain and posttranslational acylation or phosphorylation are known to play a role in membrane association [41].

Folden *et al.* [44] found that *Borrelia burgdorferi* enolase was exposed on the cell surface using microscopy and immunoassays and acted as a receptor for plasminogen. They also examined the conversion of plasminogen to plasmin and immunoblotted enolases of recombinant and cell lysates. These enolases proven to be strongly seropositive are consistent with our results.

### Elongation Factor Tu

Translational elongation factor Tu (EF-Tu) contributes to the lengthening of the peptide chain in protein synthesis [45, 46], but has additionally been reported as an adhesin, necessary for cell invasion in several pathogenic bacteria. Nieve *et al.* [47] identified AhpC, DnaK, and EF-Tu as highly immunogenic proteins of the closely related *Burkholderia thailandensis*. They found that intranasal immunization using a recombinant EF-Tu was able to generate an IgG antibody that recognized the native EF-Tu, representing a potential vaccine target for *B. pseudomallei*.

Granto *et al.* [48] identified EF-Tu in *Lactobacillus johnsonii* NCC533 (La1) as immunogenic surface molecules which mediated attachment to intestinal epithelial cells and mucins. Kolberg *et al.* [49] also found EF-Tu to be surface exposed in multiple bacteria using flow cytometry. It appears that EF-Tu is highly immunogenic in multiple species and plays a critical role for pathogens in invasion of host cells.

### LysR family transcriptional regulator

LTTRs are one of the most abundant types of transcriptional regulators in known in prokaryotics[50]. These proteins regulate diverse genes involved in virulence, metabolism, and motility. Rainbow et al. (2002) report that the proteins of the LysR family possess a potential ‘helix-turn-helix’ DNA-binding motif that is conserved in their N-terminal section.

### Sigma-54 dependent DNA binding response regulator

AtoC are regulators responsive to a variety of environmental signals. Chemical and metabolic changes modulate the expression or the activity of regulatory proteins for determining the level of expression of sigma54-dependent genes and hence the diverse bacterial functions that they encode. Sigma-54 proteins are widespread among bacteria and required for diverse functions such as motility, phage shock response, and nitrogen assimilation (Reitzer, 2003)

### Immunogenic variable proteins

Most of the immunogenic proteins are specific to serum and strain. For examples, AtpD, AtpA, DnaK, flagellum (FliC), and sucrose-6-phosphate hydrolase are common immunogenic proteins identified previously by Felgner et al. [14], but our study has shown that they are detected in not all cases. It is postulated that the variation in detected immunogenic proteins can be caused by different stages of illness and also that the different clinical presentation reflects different pathways of virulent protein involvement [29].

Gene expression studies have shown that ATP synthases such as AtpA and AtpD can be down-regulated. In host-pathogen interactions, the bacteria in the intracellular state adjust nutrient availability for survival, resulting in down regulation of energy metabolism such as ATP synthase and NADH dehydrogenase [51]. Tuanyok *et al.* [52] found that iron stress results in modulation of expression of ATP synthase subunit epsilon. But this research found strong reactivity for ATP. It is postulated that the proteins are required for surviving and thriving in the host.

FliC protein is believed to play roles of motility of the bacterium to disseminate from localized sites, such as skin and lung, to other organs via bacteremic spread. This ability of *B. pseudomallei* may be important also for intracellular survival and in the pathogenicity of both acute and chronic infection [53]. Chua et al. [54] constructed isogenic deletion mutant with flagellum gene (fliC) and challenged mice with the mutant; all animals infected with this mutant remained healthy. The patient whose serum linked to *B. pseudomallei* PB1007001 presented with a skin abscess and associated swollen regional lymph node. This illness followed a presumptive inoculating skin cut on the leg. From this patient serum, FliC was detected as an immunogenic protein.

DnaK is a chaperone protein having conserved domain like GroEL, which remains unchanged level at different media and surrounding environment and keep the transcript level even in different strains [55]. That means this chaperone protein had the sample transcript level across all the four stains at this research. At this research, two sera showed reactivity to DnaK. This protein was also identified as being immunoreactive with melioidosis case sera by others [14, 30]. Being detected by only two isolates out of four may reflect different pathway of infection or progress of infection.

## Comparison of identified common immunogenic protein sequence

We blasted our commonly detected immunogenic protein sequences with those of selected commensal bacteria, *B. mallei* and *B. thailandensis* strains to investigate structural similarities. Most of our common immunogenic proteins are chaperonins, which have highly conserved structure. Those conserved structures have high similarity with commensal bacteria and other genetically similar species. This could hinder these proteins from being used as a specific biomarkers for diagnosis or as vaccine target. Vaccine candidates should be differentiated between beneficial and harmful microbes.

The sequence comparison of the common proteins showed that the similarities were lower with commensal bacteria. These are of greatest concern as they can colonize healthy humans and are distributed widely. The lower similarities imply that the identified 8 proteins might be useful as vaccine candidates and would not cause an immune reaction to commensal microbes. The common proteins of *B. pseudomallei* matched or were very high similarity to *B. mallei* and *B. thailandensis*. However, Ndh and AtoC of *B. pseudomallei* showed the slightly lower similarity with potential as a diagnosis biomarker.

Translating our proteome data to other proteome studies of *B. pseudomallei* recovered from different clinical presentations and to serial sera responses over the time course of infections of individual patients will help further unravel the pathogenesis of infection with *B. pseudomallei.*

## Conclusion

We identified proteins common to all four strains that are strongly immunogenic with all sera used. Perhaps not coincidentally, these proteins are also highly expressed as visualized with 2DE and silver staining. This suggests that this group of eight known proteins have a potential for use as diagnostic targets or vaccine subunits. Further studies of more patient-*B. pseudomallei* isolate matches are required to assess the universality of these protein responses and to also determine how the progression of disease affects the expression of *B. pseudomallei* proteins and the resulting host antibody production.

## Acknowledgement

This work was supported by the United States Department of Homeland Security (https://www.dhs.gov/) grant no. HSHQDC-10-C-00135 and Defense Threat Reduction Agency (http://www.dtra.mil/) grant no. HDTRA1-12-C-0022. We thank our laboratory colleagues at the Royal Darwin Hospital for their support and expertise in diagnosis of melioidosis and identification of *B. pseudomallei,* and Vanessa Theobald and Glenda Harrington at the Menzies School of Health Research for laboratory assistance with this study.

